# Investigations into the emergent properties of gene-to-phenotype networks across cycles of selection: A case study of shoot branching in plants

**DOI:** 10.1101/2022.02.03.479064

**Authors:** Owen M. Powell, Francois Barbier, Kai P. Voss-Fels, Christine Beveridge, Mark Cooper

## Abstract

Predictive breeding is now widely practised in crop improvement programs and has accelerated selection response (i.e., the amount of genetic gain between breeding cycles) for complex traits. However, world food production needs to increase further to meet the demands of the growing human population. The prediction of complex traits with current methods can be inconsistent across different genetic, environmental, and agronomic management contexts because the complex relationships between genomic and phenotypic variation are not well accounted for. Therefore, developing gene-to-phenotype network models for traits that integrate the knowledge of networks from systems biology, plant and crop physiology with population genomics has been proposed to close this gap in predictive modelling. Here, we develop a gene-to-phenotype network for shoot branching, a critical developmental pathway underpinning harvestable yield for many crop species, as a case study to explore the value of developing gene-to-phenotype networks to enhance understanding of selection responses. We observed that genetic canalization is an emergent property of the complex interactions among shoot branching gene-to-phenotype network components, leading to the accumulation of cryptic genetic variation, reduced selection responses, and large variation in selection trajectories across populations. As genetic canalization is expected to be pervasive in traits, such as grain yield, that result from interactions among multiple genes, traits, environments, and agronomic management practices, the need to model traits in crop improvement programs as outcomes of gene-to-phenotype networks is highlighted as an emerging opportunity to advance our understanding of selection response and the efficiency of developing resilient crops for future climates.

## Introduction

Due to an increasingly harsh and unpredictable climate, improving the consistency and scope of predictions for crop performance is crucial for global agriculture to meet the challenge of feeding a global population of 10+ billion people (Reynolds et al., 2021). Current prediction methods used in crop breeding assume a simplified linear relationship between genotype and phenotypes (Falconer and Mackay, 1996; Meuwissen et al., 2001; Cooper et al., 2014; Walsh and Lynch, 2018; Gianola, 2021), thus limiting the realized selection response achieved by many crop improvement programs (Kholová et al., 2021). Although this simplified genotype-to-phenotype relationship (G2P map) is sufficient to successfully model the average selection trajectory of large populations (Cooper et al., 2014), this approach captures only a subset of all performance outcomes, potentially leading to misalignments between predicted performance and realized performance in the field. Such simplified G2P relationships can hinder accurate predictions for crop performance in specific management and environment combinations (Kholová et al., 2021).

Developing G2P network models that integrate the knowledge of networks from systems biology and physiology with population genomics may improve modelling of the genotype-to-phenotype relationship for many complex traits (Benfey and Mitchell-Olds, 2008; Marjoram et al., 2014; Marshall-Colon et al., 2017; Eshed and Lippman, 2019; Cooper et al., 2020). For crop improvement programs, the detected interactions in network models can unmask existing genetic variation or identify intermediate traits that can increase the selection accuracy and efficiency of developing novel crop varieties and hybrids. Applying the framework of the “breeders equation” (Falconer and Mackay, 1996; Paixão and Barton, 2016; Walsh and Lynch, 2018), the contributions of the genetic interactions (epistasis) to total genetic variation must be converted into additive genetic variation to deliver a sustainable selection response in breeding programs (Technow et al. 2021). A strong theoretical understanding exists of the importance of epistasis for selection response when based on G2P models without molecular network models (Paixão and Barton, 2016; Walsh and Lynch, 2018). However, relevant knowledge of selection response for G2P models based on networks operating over biological scales (Marshall-Colon et al., 2017; Hammer et al., 2019; Wu et al., 2019; Tardieu et al., 2020) is lagging due to the lack of adequately characterized empirically-based examples.

Here, we use axillary bud outgrowth, a primary driver of shoot branching, as a case study of an agronomically important, well-characterized and empirically described network (Bertheloot et al., 2020). The core structure of this network is largely conserved across herbaceous model plants and divergent crops. It involves different endogenous signals interacting with each other to regulate shoot branching. Hormones including auxin, cytokinins and strigolactones play a crucial role in regulating this process (Domagalska and Leyser, 2011; Barbier et al., 2019). The growing shoot apex produces auxin, which travels downwards in the stem, inhibiting cytokinin synthesis and accumulation and promoting strigolactone synthesis. Cytokinins and strigolactones induce and repress bud outgrowth, respectively. Sugar availability is a determining factor for plant growth and development, including shoot branching (Barbier et al., 2019; Fichtner et al., 2021). Axillary buds require a source of sugar to grow out, and the strong demand for sugar by the shoot apex inhibits axillary bud outgrowth (Mason et al., 2014; Barbier et al., 2019). During shoot branching, sugars were reported to promote cytokinin synthesis and inhibit strigolactone perception (Barbier et al., 2019; Bertheloot et al., 2020; Salam et al., 2020; Patil et al., 2021). Despite the detailed knowledge of molecular physiological mechanisms, translating these discoveries into breeding outcomes is still a challenge.

Therefore, the objectives of this study were threefold: (1) to extend the Bertheloot et al. (2020) model of the shoot branching network to include genetic variation for nodes of the network; (2) apply quantitative genetic methods (Falconer and Mackay, 1996; Walsh and Lynch 2018) to undertake *in silico* investigations of important G2P properties of the extended shoot branching network model that can influence breeding outcomes; and (3) develop hypotheses of the expected selection trajectories for levels of the branching network nodes and branching trait outcomes for experimental investigation. Applying the framework developed to link quantitative genetic models for trait genetic variation with crop growth models to model plant responses to environmental variation, outlined in Cooper et al. (2020), we created a shoot branching G2P network (Fig. 1) underpinned by genomic variation to demonstrate how variation from interactions in network-based G2P models is translated into selection responses for complex traits. *In-silico* selection experiments were performed on a large, segregating plant population to quantify direct (time to bud outgrowth) and indirect (intermediate traits; hormones and sucrose) selection responses. The results are discussed in terms of practical implications for developing G2P models to accelerate crop genetic improvement within established and future crop improvement programs.

**Figure 1.**
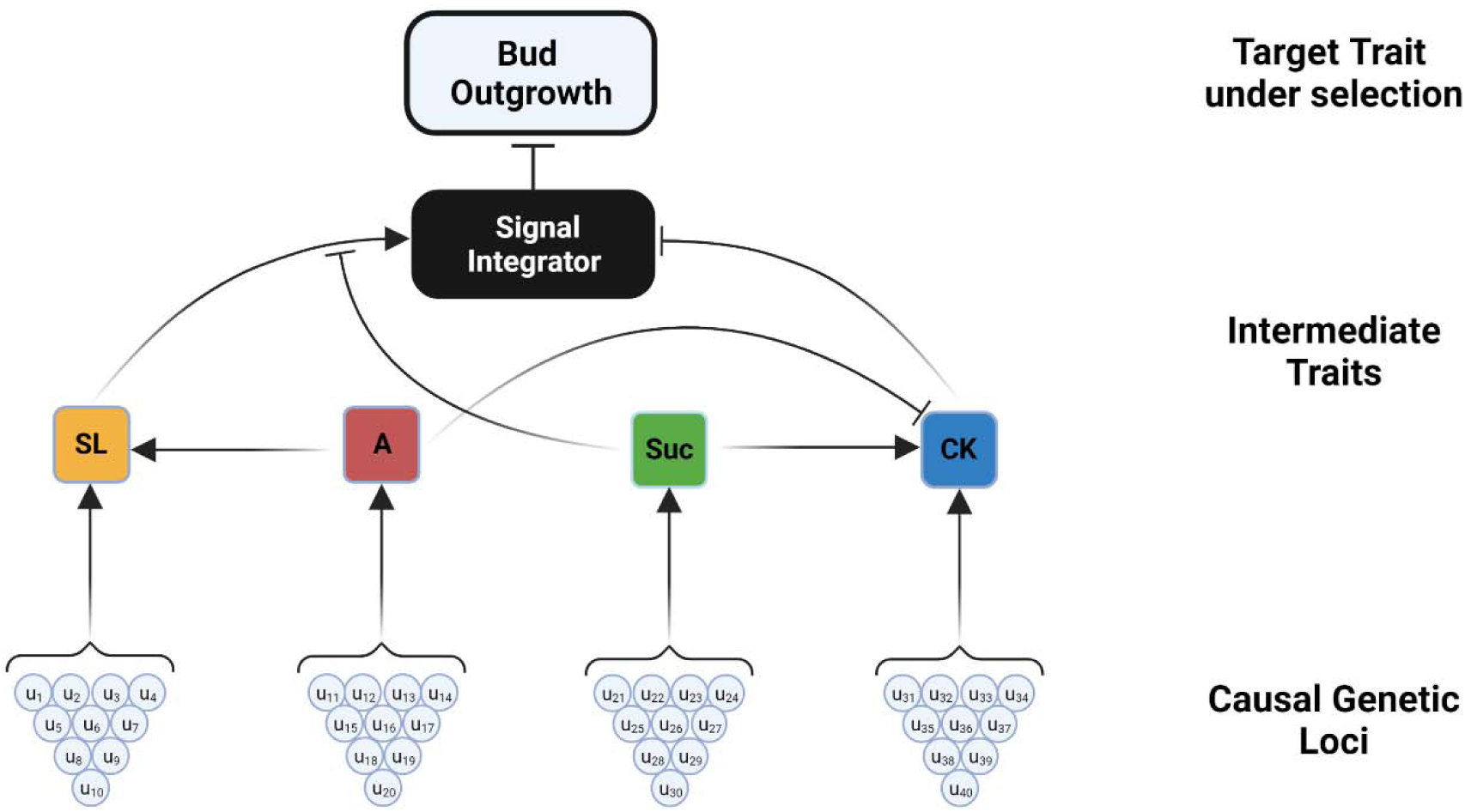
Shoot branching gene-to-phenotype (G2P) network. Ten causal genetic loci (**u**) and interactions determine the levels of each of the intermediate traits : strigolactones (**SL**), auxin (**A**), sucrose (**Suc**), and cytokinins (**CK**). In turn, levels of these intermediate traits (hormones, sucrose and the signal integrator), their interactions, and random error (**e**) determine the time to bud outgrowth of an individual plant.

## Materials and Methods

### Overview

We created a gene-to-phenotype (G2P) network model for bud outgrowth by connecting a published, empirical shoot branching network (Bertheloot et al., 2020) to underlying allelic variation in the genome. The shoot branching network models phenotypic variation for the trait “time to bud outgrowth” as the outcome of the intermediate traits, auxin, cytokinins, strigolactones, and sucrose, and their interactions (Fig.1). We simulated the additive genetic effects of 10 non-pleiotropic, causal genetic loci for each intermediate trait. The additive genetic effects of the 10 causal genetic loci determined the additive genetic values for each intermediate trait for individual genotypes.. These additive genetic values replaced the synthesis term of the intermediate traits (hormones and sucrose), in the differential equations provided by Bertheloot et al. (2020) which were used to calculate the levels for each intermediate trait (***g***). Therefore, the trait under selection, time to bud outgrowth (***y*_*BO*_**), can be viewed as a function **F** of the levels of the genotype-dependent intermediate traits (***g***) and a random error term (***e***).

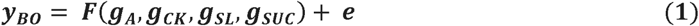

To quantify response to selection, we performed *in-silico* divergent selection experiments for increased and decreased levels of ***y*_*BO*_** over 30 selection cycles. The results presented are generated from 100 replicates of the *in-silico* selection experiment. Code for the shoot branching network, genetic simulations and figures can be accessed in the following repository: https://github.com/powellow/GeneticCanalizationOfG2PNetworks.

### Description of the Empirical Shoot Branching Model

Bertheloot et al. (2020) used experimental data to describe phenotypic variation for time to bud outgrowth within a shoot branching network as a system of differential equations. The shoot branching network takes levels of auxin and sucrose as inputs, calculates cytokinins, strigolactones and signal integrator as intermediate trait outputs, and the time to bud outgrowth (days) as the final trait output. The levels of the intermediate trait outputs are described by differential equations, which each contained three terms: (i) a synthesis term, (ii) an interaction term and (iii) a degradation term. Bertheloot et al. (2020) applied a grid search approach, using observed times to bud outgrowth and levels of cytokinins, to parameterize the coefficients of the differential equations.

### Description of the in-silico Gene-to-Phenotype Shoot Branching Network

To quantify the response to selection for time to bud outgrowth in a breeding population, a G2P shoot branching network model was developed. The G2P network connected phenotypic variation within the shoot branching network to allelic variation across a simulated genome. The simulated genomes of the individuals within a reference population of genotypes (Cooper et al., 2020) consisted of a single chromosome with 40 causal genetic loci. Each intermediate trait received additive genetic effects (***u***) from 10 non-pleiotropic causaal genetic loci. The magnitudes of ***u*** were sampled from a normal distribution, but the sum of their effects was constrained so that the additive genetic values (***a***) of individuals were within the range observed in experimental data (Table S4, Bertheloot et al. (8)). The additive genetic values (***a***)) for each intermediate trait were computed by summing the 10 additive genetic effets effects (***u*_*i*_**) according to the genotypes at the casusal geneti loci of each individual:

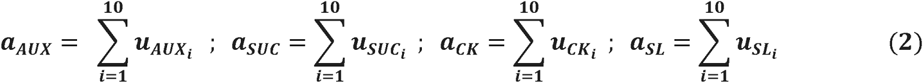

The additive genetic values of individuals replaced the synthesis terms in the differential equations, but all other steps remained unchanged from Bertheloot et al. (2020). We provide the following, adapted equations replacing the synthesis terms with the appropriate breeding values purely for thoroughness and reproducibility. The interaction terms (***γ***)and levels of intermediate traits (***g***) were calculated based on the addtive genetic values (***a***) of individuals as follows:

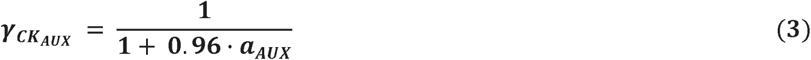

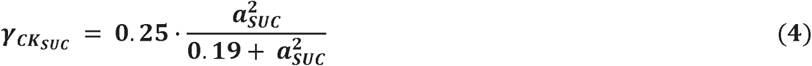

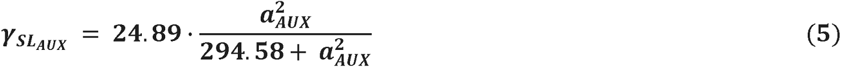

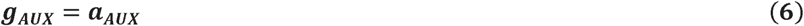

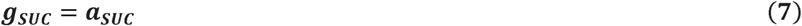

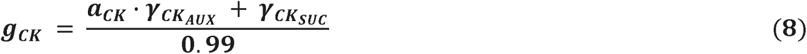

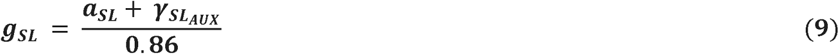

The levels of the intermediate traits were passed through a signal integrator, *I*.

Calculated as follows:

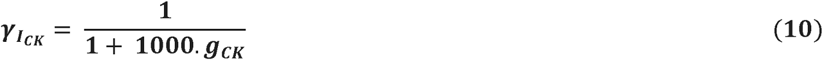

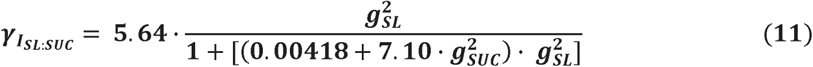

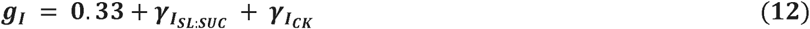

The level of the signal integrator (***g*_*I*_**) was then used to calculate the output trait, time to bud outgrowth (***g***_***BO***_), as well as the calculation of a threshold bud outgrowth trait:

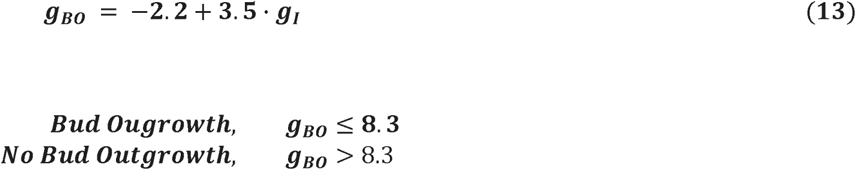

### Description of the in-silico Selection Experiments

For the *in-silico* selection experiments, we created an initial reference population of genotypes (RPG) from a single biparental cross, consisting of 1,000 F2 individuals. Phenotypes for time to bud outgrowth (***y***_***BO***_) of the 1,000 individuals of the reference population of genotypes were generated by adding a random error effect(*e* ∼ ***N***[0.*v*_*e*_]), to collectively represent stochastic environmental, developmental noise and measurement error, to the true genetic value of time to bud outgrowth, ***g***_***BO***_ (Eqn. 1). The value of ***v***_*e*_ was calculated such that the broad-sense heritability, ***H***^2^ of time to bud outgrowth ranged between 0.1 and 1.0 in the initial RPG. We present results for ***H***^2^ = 0.3 and 1.0. The value of ***v***_*e*_ was held constant over the 30 selection cycles. Therefore, individuals experienced the same level of random error throughout the experiment, while the ***H***^2^ for time to bud outgrowth could change with the magnitude of genetic variance in the reference population of genotypes due to selection. The population underwent 30 cycles of truncation selection with discrete, non-overlapping generations (cycles) for either higher or lower time to bud outgrowth. For each cycle, the ‘best’ 100 individuals were selected based on their output trait phenotype, time to bud outgrowth, to be used as parents (selection proportion = 0.1) and crossed at random to create 1,000 offspring for the next cycle of evaluation and selection. Independent population replicates were generated by repeating the whole *in-silico* selection experiment process one hundred times.

### Hierarchical Clustering of Selection Trajectories

Multi-trait performance landscapes for the intermediate trait phenotypes were generated to investigate quantitative genetic properties of the selection trajectories over 30 cycles of selection (Gavrilets, 2004; Messina et al., 2011; Walsh and Lynch, 2018; Cooper et al., 2020). To aid the visualization of the exploration of the shoot branching performance landscape via selection, the selection trajectories of the 100 replicates underwent hierarchical clustering. The 100 replicates were classified into 3 clusters using Ward ’s method (Ward, 1963; Wishart, 1969; Williams, 1976). Rows of the matrix corresponded to the replicate id for each of the 100 replicates, columns of the matrix corresponded to each of the 30 selection cycles, and the cells contained values for strigolactone levels. The group mean trait levels for each selection cycle were plotted on the performance landscapes to visualize the selection trajectories.

## Results

The *in-silico* selection experiments with the shoot branching G2P network (Fig. 1) revealed the presence of cryptic genetic variation for the intermediate traits, hormones, and sugars that the selection for time to bud outgrowth struggled to access (Waddington, 1942; Flatt, 2005; Masel, 2006; Walsh and Lynch, 2018). Such cryptic sources of genetic variation occur when selection cannot directly translate the sources of genetic variation into a selection response to improve adaptation and performance. The cryptic genetic variation within the *in-silico* experiment led to many repeated selection cycles with reduced selection response and large variation in selection trajectories across different replicate populations (Fig. 2*A, C*). Despite the large magnitudes of cryptic genetic variation for the intermediate traits under indirect selection, only small differences were observed among the selection trajectories of time to bud outgrowth, the output trait under direct selection.

**Figure 2.**
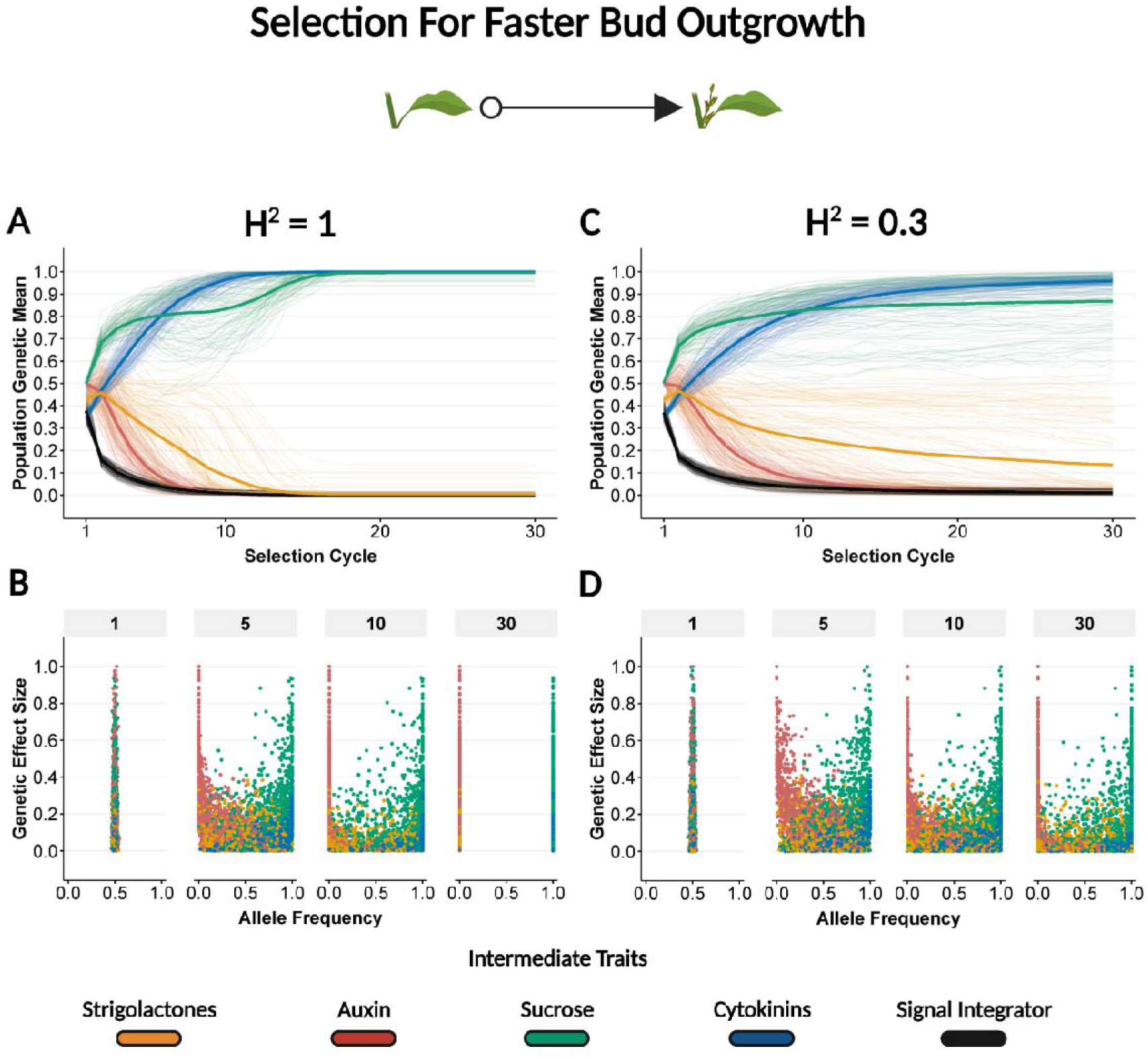
Selection trajectories for the shoot branching G2P network for faster bud outgrowth and the relationships between normalized genetic effect sizes and allele frequency changes at causal genetic loci over selection cycles. ***(A, B)*** Results from selection with a broad sense heritability,, of 1. ***(C, D)*** Results from selection with a broad sense heritability,, of 0.3. ***(A, C)*** Normalized total genetic values for the intermediate traits over selection cycles. Thick lines are the normalized genetic means averaged across the population replicates. Thin lines are the normalized genetic means for each population replicate. ***(B, D)*** Plot of allele frequency changes at causal genetic loci versus normalized genetic effect sizes. Each point represents a causal genetic locus from the 100 population replicates. Values are presented for selection cycles 1, 5, 10, and 30.

Cryptic genetic variation for the intermediate traits resulted in temporary and permanent plateaus in selection response (Fig. 2*A, C*). The emergence of these plateaus began after only three selection cycles. In scenarios with a simulated broad-sense heritability of 1 (no stochastic error variation during selection), the average genetic mean for sucrose began to increase again around selection cycle nine, when the genetic mean for strigolactones reached approximately 0.1. However, even with this perfect selection accuracy, a few of the replicate populations reached permanent plateaus at local maxima for two intermediate traits, sucrose and strigolactones. In scenarios with error included in the phenotypes of time to bud outgrowth, selection response was decreased for all shoot branching G2P network components, with the largest reductions observed for the genetic mean of sucrose. For example, with a broad-sense heritability of 0.3, the average genetic mean for sucrose across the 100 population replicates reached a permanent plateau at less than 90% of the maximum theoretical value after 30 selection cycles (Fig. 2*C*), with several individual populations achieving less than 60%.

Different magnitudes of cryptic genetic variation across the intermediate traits of hormones and sucrose resulted in different allele frequency changes for causal loci with similar genetic effect sizes for the intermediate traits (Fig. 2*B, D*). This trend was most apparent for causal loci occupying the bottom 60% of genetic effect sizes. In this G2P network simulation, alleles of causal loci with moderate genetic effect sizes (0.2–0.6) for cytokinins and auxin reached fixation within the reference population of genotypes (allele frequency of 0 or 1). However, causal loci for sucrose and strigolactones, also with moderate genetic effect sizes, were still segregating in the reference population of genotypes after 30 selection cycles (Fig. 2*D*). In the most extreme cases of the population replicates, causal loci with small genetic effect sizes (<0.2) underwent genetic drift in the reference population of genotypes, with allele frequency changes in the opposite direction from that expected based on the direction of selection. For example, we observed increases in allele frequencies of causal loci for strigolactones until at least selection cycle 10 (Fig. 2*B, D*) even though, in the absence of genetic drift or interaction effects, these would be expected to be selected against under the direct selection for faster bud outgrowth (Fig. 1). This property of intermediate traits was independent of the heritability of the trait under direct selection.

Performance landscapes were generated to further investigate and visualize emergent properties at the intermediate levels of the shoot branching network (Fig. 3). Steep gradients for time to bud outgrowth were observed at intermediate values of the intermediate traits, strigolactones and sucrose. Flatter gradients for time to bud outgrowth values were observed at extreme values of the intermediate traits. The plateaus in the performance landscape reflect that the large sources of genetic variation for sucrose and strigolactones translated into only a small variation in values of time to bud outgrowth. The average selection trajectories of the population replicates followed the steepness of performance landscapes, with a consistent higher strength of selection for higher sucrose levels in the first few selection cycles, followed by selection for lower strigolactone levels (Fig 3A & 3B). Although there was considerable variability in the selection trajectories among the individual population replicates (Supplementary Fig. 1). The inclusion of stochastic error in time to bud outgrowth values, ***H***^2^ = 0.3, resulted in selection trajectories stopping at a local maximum instead of the global maximum (Fig. 3B).

**Figure 3.**
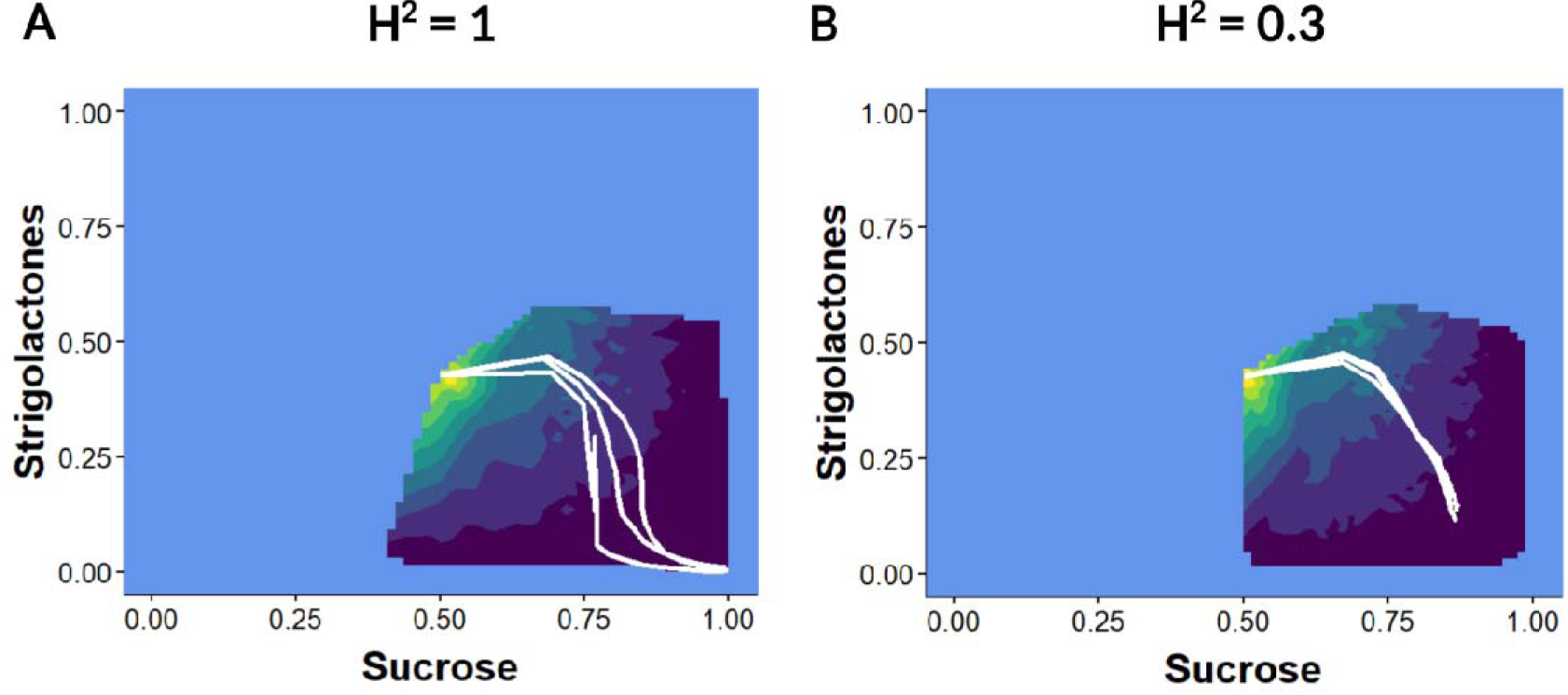
Exploration of the Shoot Branching Performance Landscape via Selection. ***(A)*** Results from selection with a broad sense heritability,, of 1. ***(B)*** Results from selection with a broad sense heritability,, of 0.3. To aid visualization, the selection trajectories of 100 population replicates were grouped into 3 clusters using Ward ‘s method (see Methods). The average selection trajectories of the three clusters (thick white lines) are plotted against the normalised total genetic values for strigolactones and sucrose. The contour shows the variation in strigolactone and sucrose values and the values for time to bud outgrowth observed across the 100 population replicates. The light blue area depicts the area of the full shoot branching performance landscape, determined by the Bertheloot et al. (2020) model, unexplored by selection.

## Discussion

In this study, cryptic genetic variation (Waddington, 1942; Masel, 2006; Walsh and Lynch, 2018) accumulated over selection cycles for the intermediate traits (hormones and sucrose) of the shoot branching G2P network, which resulted in reduced selection responses. The cryptic genetic variation for sucrose can be explained by the complex interaction between sucrose and strigolactone signalling in the G2P network (Fig. 1), resulting in genotypes with completely different combinations of strigolactone and sucrose levels producing similar values for time to bud outgrowth (Fig. 4). The occurrence of multiple intermediate G2P states mapping to fewer output trait states, associated with the emergent quantitative genetic property of cryptic genetic variation identified for the branching network model (Fig. 1), complicates the prediction of phenotype from genotype and the outcomes of selection strategies (Fig. 2 & 3), as implemented in plant breeding programs.

**Figure 4.**
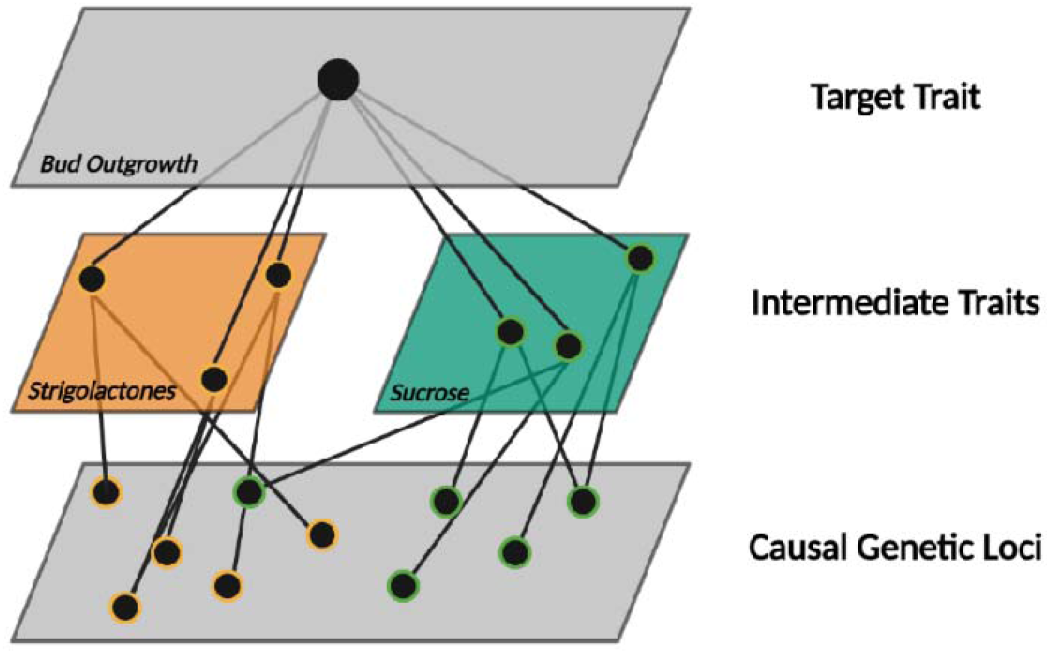
Genetic Canalization of Gene-To-Phenotype Networks. Multiple genetic combinations of intermediate traits produce similar values for the target trait, causing the accumulation of cryptic genetic variation that cannot be accessed by selection.

The accumulation of cryptic genetic variation is not specific to the shoot branching G2P network. It can occur whenever non-linear relationships exist among traits or casual genetic loci due to genetic canalization (Waddington, 1942; Flatt, 2005). Therefore, we expect genetic canalization to be pervasive in complex traits under selection that result from interactions among multiple interacting genes (Kauffman et al., 2004; Rünneburger and Le Rouzic, 2016; Ødegård and Meuwissen, 2016), traits, environments, and agronomic management practices. The expectation of the emergent property of genetic canalization for G2P networks controlling complex traits raises several important questions for crop improvement programs.

What are the impacts of reductions in detectable genetic variation for complex traits under selection? Reductions in detectable genetic variation, in part due to genetic canalization, have important implications for the selection responses achieved by crop improvement programs. Reductions in detectable genetic variation for complex target traits reduce the accuracy of identifying the best performing individuals, which leads to plateaus in selection response (Fig. 2A, 2C, 3B). Of even more concern, the combination of reduced accuracy and reduced detectable genetic variation could result in mistaking plateaus associated with local maxima for true, permanent selection limits for breeding populations (Fig. 3B).

The results obtained from the *in-silico* selection experiment based on the branching network (Fig. 1) highlight an important G2P prediction question for plant breeders; How can crop improvement programs promote decanalization? Decanalization would allow the release of cryptic genetic variation for complex traits and accelerate selection response for the reference population of genotypes. In this study, decanalization and the subsequent increases in selection response occurred via the fixation of causal loci by chance (genetic drift) at different cycles for different population replicates (Fig 2B, D). Similar decanalization events could be a contributing factor to the unexpected (according to an additive finite locus model and the breeder’s equation; (Falconer and Mackay, 1996; Walsh and Lynch, 2018)), continued selection responses seen in long-term selection experiments (Dudley, 2007; Goodnight, 2015). A more targeted strategy would be to restructure crop breeding programs to promote and control the conversion of non-additive, epistatic genetic variance into additive genetic variance within the reference population of genotypes (Cooper et al., 2005). For example, in maize (*Zea mays*), Technow et al. (2021) demonstrated that a decentralized structure of multiple, smaller crop improvement programs interconnected by a few key parents was required to facilitate selection response under high levels of G2P genetic complexity. Another complementary strategy involves direct selection on traits at intermediate layers of the G2P network to circumvent the complex interactions that generate canalization of traits at higher levels of the network hierarchy. In the case of shoot branching, decanalization could be achieved by direct selection on sucrose or strigolactone levels. The many influences of such emergent non-linear properties and the need to consider alternative breeding strategies to accelerate improvement of the complex traits motivate experimental-simulation investigations to develop appropriate G2P models (Marshall-Colon et al., 2017; Hammer et al., 2019; Cooper et al., 2020; Tardieu et al., 2020)

The development of G2P networks that link, empirically based, plant models with natural genomic variation is crucial to appropriately design strategies and experiments to tackle many compelling questions in crop improvement. This study is a demonstration of the ideas of modelling selection response through crop growth models outlined in Cooper et al. (2020), using the plant branching network model developed by Bertheloot et al. (2020). A benefit of taking such a view of the G2P relationship was the observation of genetic canalization as an emergent property of selection for faster shoot branching, which would not have been achievable taking standard physiological modelling or single-trait quantitative genetics approaches in isolation. The multi-trait structure and interactions within plant models generate conditional effects that contribute to the quantitative genetic properties of epistasis and vertical pleiotropy when connected to genomic variation. Specifying the genetic effects of the inputs of plant models at the level of genes instead of genotypes allows the assessment of selection response over multiple selection cycles, as is required for the design of breeding strategies (Hammer et al., 2019; Cooper et al., 2020). Such G2P networks also include genetic constraints enforced by processes such as recombination and linkage to provide more realistic predictions of the exploration of performance landscapes of traits (Messina et al., 2011; Technow et al., 2021). In this study, selection for faster shoot branching explored a relatively small area of the total performance landscape encoded by the Bertheloot et al. (2020) branching model (Fig. 3).

Future studies can exploit the increased power and flexibility when viewing complex traits as gene-to-phenotype networks in: (i) *in silico* simulations (Hammer et al., 2019; Cooper et al., 2020), akin to our approach; (ii) empirical, longitudinal, “select and sequence” studies (Lenski and Travisano, 1994; Wisser et al., 2019) or (iii) broader exploration of performance landscapes with genome editing of network components (Eshed and Lippman, 2019) to improve understanding of selection response of complex traits in nature and agriculture.

## Acknowledgements

We thank Dr Franziska Fichtner for fruitful discussions about the molecular physiology of the shoot branching network and comments on the manuscript. Figures were created or formatted with Biorender.

## Author Contributions

OMP and MC conceived the study and helped interpret the results. OMP developed the gene-to-phenotype network, developed the in-silico selection experiments, performed the analyses, and wrote the manuscript. FB provided input on the shoot branching model, helped interpret the results, and wrote parts of the introduction. KVF developed a previous iteration of the G2P network. CB provided input on the shoot branching model and helped interpret the results. All authors read, refined, and approved the final manuscript.

---

**Supplementary Figure 1.**
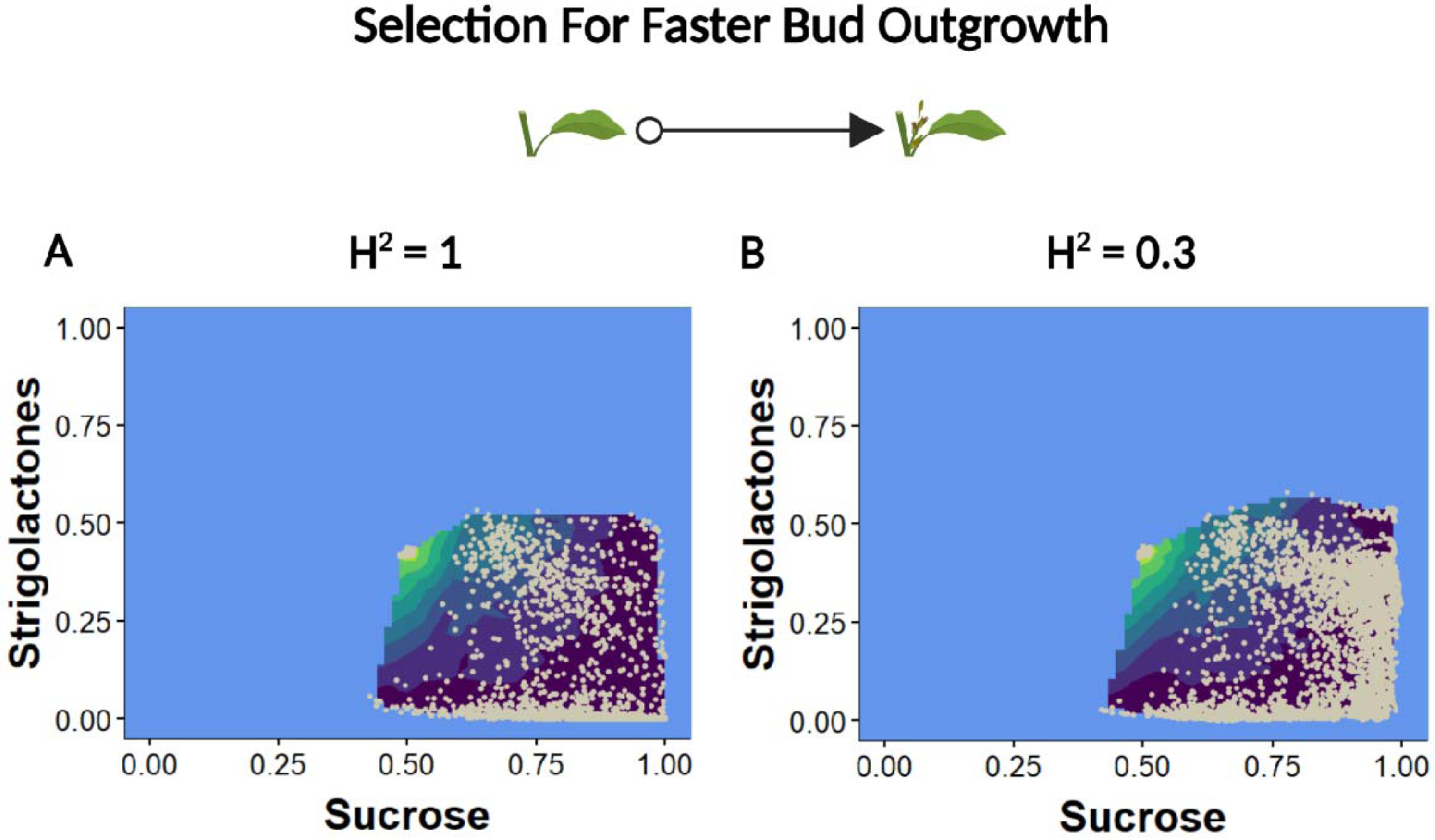
Population variation during the exploration of the shoot branching performance landscape. ***(A)*** Results from selection with a broad sense heritability,, of 1. ***(B)*** Results from selection with a broad sense heritability,, of 0.3. The contour shows the variation in strigolactone and sucrose values and the values for time to bud outgrowth observed across the 100 population replicates. The light blue area depicts the area of the full shoot branching performance landscape, determined by the Bertheloot et al. (2020) model, unexplored by selection. Grey dots show average strigolactone and sucrose values for population replicates at a particular selection cycles.

